# Does training method matter?: Evidence for the negative impact of aversive-based methods on companion dog welfare

**DOI:** 10.1101/823427

**Authors:** Ana Catarina Vieira de Castro, Danielle Fuchs, Stefania Pastur, Liliana de Sousa, I Anna S Olsson

## Abstract

There is a growing number of dogs kept as companion animals, and the methods by which they are trained range broadly from those using mostly positive punishment and negative reinforcement (aversive-based methods) to those using primarily positive reinforcement (reward-based methods). Although the use of aversive-based methods has been strongly criticized for negatively affecting dog welfare, these claims do not find support in solid scientific evidence. Previous research on the subject lacks companion dog-focused research, investigation of the entire range of aversive-based techniques (beyond shock-collars), objective measures of welfare, and long-term welfare studies. The aim of the present study was to perform a comprehensive evaluation of the short- and long-term effects of aversive- and reward-based training methods on companion dog welfare. Ninety-two companion dogs were recruited from three reward-based (Group Reward, n=42) and four aversive-based (Group Aversive, n=50) dog training schools. For the short-term welfare assessment, dogs were video recorded for three training sessions and six saliva samples were collected, three at home (baseline levels) and three after the training sessions (post-training levels). Video recordings were then used to examine the frequency of stress-related behaviors (e.g., lip lick, yawn) and the overall behavioral state of the dog (e.g., tense, relaxed), and saliva samples were analyzed for cortisol concentration. For the long-term welfare assessment, dogs performed a cognitive bias task. Dogs from Group Aversive displayed more stress-related behaviors, spent more time in tense and low behavioral states and more time panting during the training sessions, showed higher elevations in cortisol levels after training and were more ‘pessimistic’ in the cognitive bias task than dogs from Group Reward. These findings indicate that the use of aversive-based methods compromises the welfare of companion dogs in both the short- and the long-term.

## 1. Introduction

To fulfil their increasingly important role as companion animals, dogs need to be trained to behave in a manner appropriate for human households. This includes, for example, learning to eliminate outdoors or walk calmly on a lead [1,2]. The fact that dog behavior problems are the most frequently cited reason for rehoming or relinquishment of dogs to shelters and for euthanasia [2] suggests that such training is often missing or unsuccessful.

Dog training most often involves the use of operant conditioning principles, and dog training methods can be classified according to the principles they implement: aversive-based methods use positive punishment and negative reinforcement and reward-based methods rely on positive reinforcement and negative punishment [3]. There is a heated debate surrounding the use of aversive-based training methods, as studies have linked them to compromised dog welfare (e.g., [4–9]). Some aversive-based tools, such as shock collars, have indeed been legally banned in some countries [10]. However, a recent literature review by [3] concluded that, because of important limitations, existing studies on the topic do not provide adequate data for drawing firm conclusions. Specifically, the authors reported that a considerable proportion of the studies relied upon surveys rather than on objective measures of both training methods and welfare; that they focused on sub-populations of police and laboratory dogs which only represent a small portion of dogs undergoing training; and, finally, that the empirical studies have concentrated mainly on the effects of shock-collar training, which is only one of several tools used in aversive-based training. In summary, limited scientific evidence exists on the effects of the entire range of dog training techniques on companion dog welfare.

Furthermore, previous empirical studies have focused on the short-term effects of training methods on dog welfare. Behavioral and physiological indicators of welfare, such as the frequency of stress-related behaviors and the concentration of salivary cortisol, have been collected in and around the training situation (e.g., [8, 11]; see also [3]). However, the long-term welfare implications of training methods have not yet been examined. To our knowledge, only one study evaluated the long-term effects of training on welfare. Christiansen et al (2001) [12] found no effect of shock collar training on dog fear or anxiety; however this was based on dog owner reports of behavior and temperament tests rather than on objective and animal-based welfare indicators. Importantly, a suitable assessment of the effects of training methods on dog welfare should comprise an evaluation of both their short- and long-term effects.

Long-term (or chronic) stress can arise from the cumulative exposure to aversive experiences [13], which may reflect the experience of dogs trained with aversive-based methods. A body of research has shown that long-term stress is associated with changes in the long-term affective state of animals (e.g., [14–16]). One way to assess affective states is through the cognitive bias paradigm (e.g., [16]). The cognitive bias task has been validated as an effective tool to evaluate the affective states of non-human animals and has been extensively used with several species, including dogs [17–19]. The rationale behind the paradigm is based on theoretical and empirical findings that an individual’s underlying affective state biases its decision-making and, specifically, that individuals experiencing negative emotional states make more ‘pessimistic’ judgements about ambiguous stimuli than individuals experiencing more positive emotional states [16, 18].

Therefore, the aim of the present study was to perform a comprehensive evaluation of the short- and long-term effects of aversive- and reward-based training methods on companion dog welfare. By performing an objective assessment of training methods (through the direct observation of training sessions) and by using objective measures of welfare (behavioral and physiological data to assess short-term effects, and a cognitive bias task to assess long-term effects), we addressed the question of whether aversive-based methods actually compromise the well-being of companion dogs. We hypothesized that dogs trained using aversive-based methods would display higher levels of stress during training, as determined by behavioral and physiological indicators of stress during training sessions, and more ‘pessimistic’ judgments of ambiguous stimuli during a cognitive bias task performed outside the training context.

Understanding the effects of training methods on companion dog welfare has important consequences for both dogs and humans. Both determining and applying those training methods that are less stressful for dogs is a key factor to ensure adequate dog welfare and to capitalize on the human benefits derived from interactions with dogs [20, 21].

## 2. Materials and methods

### 2.1. Ethical Statement

All procedures were approved by ICBAS (Abel Salazar Biomedical Sciences Institute) ORBEA (Animal Welfare Body). All head trainers of dog training schools and owners completed a consent form authorizing the collection and use of the data.

### 2.2. Training schools

#### 2.2.1. Recruitment

The first author (ACVC) searched on the internet for dog training schools in the metropolitan area of Porto, Portugal. Based on geographical proximity and on the listed training methods, ACVC posteriorly contacted eight schools through telephone. She approached the head trainers of the different schools on their willingness to participate in a study that aimed to evaluate dog stress and welfare in the context of training and explained the entire methodology. However, it was not directly revealed that the aim of the study was to compare the effects of different training methods. Seven out of the eight dog training schools agreed to participate.

The training schools had different class structures and training sites; however, the types of behaviors trained was fairly standard across training schools. These characteristics are described in detail in Appendix S1.

#### 2.2.2. Classification of training methods

After securing seven participating schools, we performed an objective assessment of the training methods used by each school. We videotaped four training sessions at each training school using a video camera on a tripod. Afterwards, we analyzed the videos for the frequency of aversive-based operant conditioning procedures utilized, namely positive punishment and negative reinforcement (see Appendix S1 for the specific definitions). The analysis was performed by AVCV using The Observer XT software, version 10.1 (Noldus Information Technology Inc, Wageningen, The Netherlands). The schools were classified as aversive-based if they used any positive punishment and/or negative reinforcement training techniques, and as reward-based if they did not use any of these techniques. Schools A, C, D and F were classified as aversive-based, and Schools B, E and G were classified as reward-based (see Appendix S1).

### 2.3. Subjects

The head trainer of each training school was asked to indicate at least fourteen dogs fitting our inclusion criteria, and we then approached the owners to ask if they were willing to participate. The information about the study given to the owners was the same that was given to the head trainers of the schools. The inclusion criteria for the dogs were: 1) to have attended the training school for less than two months, in order to mitigate habituation to training methods, and 2) to be free of behavioral problems (e.g., aggression, fearfulness and separation anxiety, as determined by the owner and ACVC), in order to prevent any confounding stress.

Over the course of the study, which was conducted between October 2016 and March 2019, the owners of 122 companion dogs agreed to participate. However, 30 dog owners dropped out of the training schools before any meaningful data could be collected. Specifically, these subjects dropped out before meeting our requirement that at least two training sessions were video recorded and that the owner completed a written questionnaire. Hence, our final sample comprised 92 subjects, 50 recruited from aversive-based schools, hereafter ‘Group Aversive’ (Schools A, D and F: 14 dogs, School C: 8 dogs), and 42 from reward-based schools, hereafter ‘Group Reward’ (School B and G: 15 dogs, School E: 12 dogs).

As for subjects’ demographics, the average age was 11.9 (±9.3) months, 54% were male and 35% were neutered/spayed. Thirty-four percent were mixed-breed dogs and the remaining 66% belonged to a FCI-recognized breed group: 18% belonged to Group 1: Sheepdogs and Cattledogs (except Swiss Cattledogs), 13% to Group 2: Pinscher and Schnauzer – Molossoid and Swiss Mountain and Cattledogs, 5% to Group 3: Terriers, 4% to Group 6: Scent hounds and related breeds, 2% to Group 7: Pointing dogs, 20% to Group 8: Retrievers – Flushing Dogs – Water Dogs, and 3% to Group 9: Companion and Toy Dogs.

### 2.4. Data collection

The study had two phases. The goal of Phase 1 was to evaluate the welfare of dogs within the training context, and the goal of Phase 2 was to evaluate the welfare of these same dogs outside the training context. These aimed to represent measures of the short- and long-term impact of training methods on the welfare of dogs.

#### 2.4.1. Phase 1 – Evaluating welfare within the training context

In order to evaluate behavioral indicators of welfare during training, each dog was videotaped for the first 15 minutes of three training sessions using a video camera on a tripod (one Sony Handycam HDR-CX405 and two Sony Handycam DCR-HC23). Five experimenters were responsible for data collection (ACVC, Danielle Fuchs – DF, Stefania Pastur – SP, and two undergraduate students, Margarida Lencastre and Flávia Canastra). The cameras were positioned to get an optimal view of the specific participant without interfering with training. The day and time of the training sessions were determined by the training schools and by the participants’ availability.

To obtain physiological data on stress during training, six saliva samples were collected per dog to allow assay of salivary cortisol [8, 22]. Three samples were collected 20 min after each training session (PT – post-training samples) and three were collected at home on days when no training took place, approximately at the same time as PT samples (BL – baseline samples). Owners were asked not to give their dog water in the 20 minutes preceding each sample collection, nor a full meal in the hour preceding each sample collection, respectively. The timing for sample collections, as well as other recommendations regarding saliva collection for cortisol analysis, were drawn from previous relevant research on dogs’ cortisol responses to potentially stressful stimuli [8, 22–24]. ACVC collected the first sample of every subject (PT1) while simultaneously demonstrating proper sample collection to the owners. The following samples were always collected by the owners. A synthetic swab (Salivette®) was rubbed in the dogs’ mouth for about 2 minutes to collect saliva. For samples collected at the training schools (PT), the swab was placed back into the provided plastic tube and immediately stored on ice. It was then transferred to a −20°C freezer as soon as possible. For samples collected at home (BL), owners were instructed to place the swab back into the plastic tube and immediately store it in their home freezer. Owners were provided with ice-cube plastic makers to transport the BL samples to the training school during the next scheduled training session without them unfreezing, and they were stored at −20°C as soon as possible. Owners were also provided with detailed written instructions for saliva collection and ACVC’s cell phone number in case any owners had questions related to sample collection. For standardization purposes, we ensured that Phase 1 did not last more than three months for each dog.

#### 2.4.2. Phase 2 - Evaluating welfare outside the training context

After finishing data collection for Phase 1, dogs moved to Phase 2, which consisted of a spatial cognitive bias task. For standardization purposes, we ensured that 1) dogs had attended the training school for at least one month prior to moving to Phase 2 and that 2) the cognitive bias task was conducted within one month of completing Phase 1. Due to limited owner availability, 13 subjects either dropped out of the study or did not meet the criteria for Phase 2, resulting in 79 (44 from Group Aversive and 35 from Group Reward) of the original 92 dogs participating in Phase 2. The tests were scheduled according to owners’ availability, both on weekdays and Saturdays.

The test was conducted in an indoor room (7.7 × 3 meters) within a research building at the Abel Salazar Biomedical Sciences Institute (ICBAS), University of Porto in Portugal. All dogs were unfamiliar with the room prior to testing. Two experimenters conducted the test while the dog’s owner(s) sat in a chair in a corner area of the room (see Figure 1). Dog owners were asked not to look into the dog’s eyes or to speak to the dog during the test, unless the experimenters instructed otherwise. The entire test took place over one meeting for each dog. The room was cleaned with water and liquid detergent at the end of each test.

**Figure 1.**
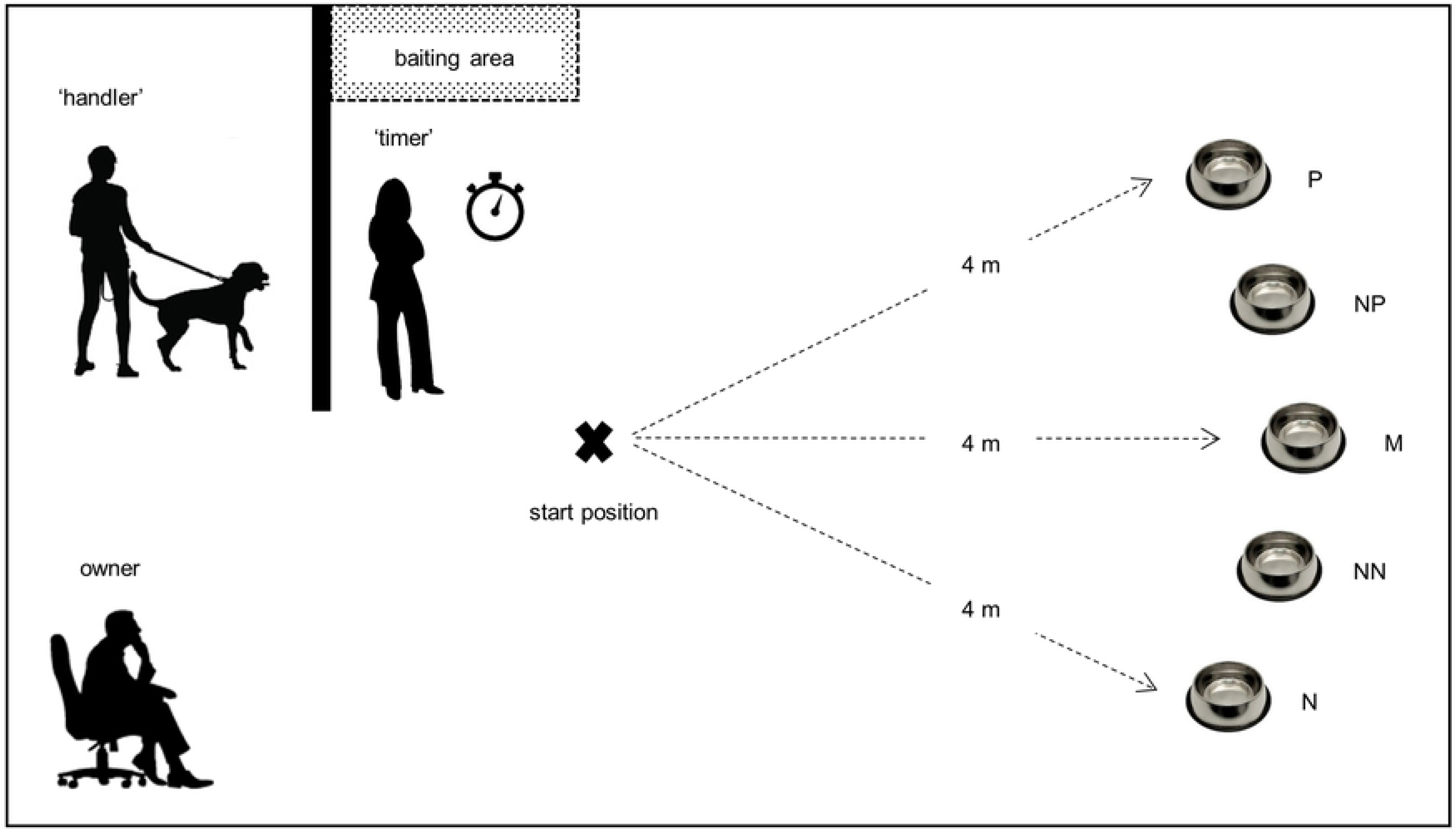
Schematic representation of the cognitive bias task.

##### 2.4.2.1. Familiarization period

Prior to the start of the cognitive bias task, the dogs were given the opportunity to familiarize with the test room and the researchers. This consisted of a 10-min period during which the dog was allowed to freely explore the room and engage with the researchers and the owner(s).

##### 2.4.2.2. Training phase

The methodology of Phase 2 was based on [19]. During the training phase, dogs were trained to discriminate between a ‘positive’ (P) location of a food bowl, which always contained a food reward, and a ‘negative’ (N) location, which never contained a food reward. At the start of each trial, the dog was held by the ‘handler’ (played by SP, DF, Margarida Lencastre, Flávia Canastra or Joana Guilherme-Fernandes) behind a barrier (2 × 2 m, see Figure 1), while the ‘timer’ (ACVC) baited (or did not bait, depending on the type of trial) the bowl with a piece of sausage (approximately 1.25 g for smaller dogs and 2.5 g for larger dogs). To ensure that the dog, the owner and the ‘handler’ were blind to whether or not the bowl contained food during each trial, the bowl was baited out of their sight, on the opposite side of the barrier. Additionally, the food reward was rubbed onto the food bowl before every trial to prevent the influence of olfactory cues. The height of the food bowl was such that visual cues to the presence or absence of food could not be judged by the dog at the start position.

After baiting (or not baiting) the bowl, the ‘timer’ placed it at one of the two training locations. The ‘timer’ then determined the start of the trial, by verbally signaling to the ‘handler’, upon which the ‘handler’ led the dog to the start position and released him. The ‘handler’ always led the dog to the start position on her left side. Because we found that dogs had some difficulty noticing the bowl at the end of the room during pilot tests, the ‘handler’ walked towards the bowl and pointed it out to the dog in the first four trials. For the remaining trials, the ‘handler’ simply walked the dog to the start position and released him. After the dog reached the food bowl and (when applicable) ate the reward, the ‘handler’ collected him and led him behind the barrier to start the next trial. The latency to reach the bowl, defined as the time elapsed between release at the start position and the dog putting his head in line with the edge of the bowl, was recorded for each trial by the ‘timer’ using a stopwatch.

The position of the ‘positive’ and ‘negative’ locations was counterbalanced across subjects and training schools, such that for half of the dogs from each training school, the ‘positive’ location was on the right hand side as they faced the test area, and for the other half it was on the left. Initially, each dog received two consecutive ‘positive’ trials (bowl placed in the ‘positive’ location) followed by two ‘negative’ trials (bowl placed in the ‘negative’ location). Subsequently, ‘positive’ and ‘negative’ trials were presented in a pseudorandom order, with no more than two trials of the same type being presented consecutively.

All dogs received a minimum of 15 training trials to learn the discrimination between bowl locations. Dogs were considered to have learnt an association between bowl location and food (the learning criterion) when, after a minimum of 15 trials, the longest latency to reach the ‘positive’ location was shorter than any of the latencies to reach the ‘negative’ location for the preceding three ‘positive’ trials and the preceding three ‘negative’ trials. Each trial lasted a maximum of 20 seconds. If the dog did not reach the bowl by that time, the trial automatically ended and a latency of 20 seconds was recorded.

All but two dogs were able to complete the training phase. For the two dogs that failed to complete training, one did not show any interest in the food reward and the other was food-motivated but could not focus on the task. These two dogs belonged to Group Aversive. Therefore, the total number of subjects completing Phase 2 in Group Aversive was 42.

##### 2.4.2.3. Test phase

Testing began once the learning criterion was achieved. Test trials were identical to training trials except that the bowl (empty) was placed at one of three ambiguous locations equally spaced along an arc 4 m from the dog’s start position, between the ‘positive’ and ‘negative’ locations. The three test locations were: ‘near-positive’ (NP: one third of the way along the arc from the ‘positive’ location), ‘middle’ (M: half way along the arc), ‘near-negative’ (NN: one third of the way along the arc from the ‘negative’ location). Three test trials were presented at each test location (nine test trials in total) in the following order for all dogs: M,NP,NN,NP,NN,M,NN,M,NP (each location was presented first, second or third in each block of three test trials). Each test trial was separated from the next one by two training trials identical to those conducted in the training phase (one ‘positive’ and one ‘negative’ trials presented in a random order), in order to maintain the associations between the ‘positive’ and ‘negative’ locations and the presence or absence of food, respectively. Thus, the test phase included a further sixteen training trials interspersed in blocks of two between the nine test trials.

To end the test phase, a final trial was conducted by placing an empty bowl in the ‘positive’ location to determine whether dogs ran to the empty bowl as quickly as they did to the baited bowl. This was meant to establish that the dogs were not relying on olfactory or visual cues during the test. During the entire test, each trial was kept as similar as possible in terms of preparation time and activity, and dogs were handled in the same way throughout the test.

Due to circumstances beyond our control, namely people speaking loudly and other dogs barking in the building during some of the tests, some subjects were clearly distracted and disengaged from the task during some trials. Whenever this happened, no latency was recorded for that trial. The experimenters waited for the dog to resettle and moved to the following trial.

### 2.5. Questionnaire

All owners were asked to complete a brief written questionnaire regarding dog demographics and background, and owner demographics and experience with dogs and dog training. The questionnaire was based on [9].

### 2.6. Data analysis

#### 2.6.1. Phase 1 – Evaluating welfare within the training context

##### 2.6.1.1. Behavior coding

We developed two ethograms based on previous literature to record the frequency of different stress-related behaviors and the time spent in different behavioral states and panting during the training sessions [8, 9, 25]. The behaviors and their definitions are described in Appendix S2.

Behavior coding was conducted by three observers (ACVC, DF and SP). DF and SP were blind to how the training schools had been classified according to training methods. Each video was coded twice, once with the ethogram for stress-related behaviors, using a continuous sampling technique (by ACVC and DF, see Appendix S2), and a second time with the ethogram for overall behavioral state and panting, by scan-sampling at 1 minute intervals (by ACVC and SP, see Appendix S2). The Observer XT software, version 10.1 (Noldus Information Technology Inc, Wageningen, The Netherlands) was used to code for stress-related behaviors and Windows Movie Player and Microsoft Excel to code for overall behavioral state and panting.

Before coding independently, observers DF and SP were trained to become familiar with the ethograms, and inter-observer reliability was assessed. Inter-observer reliability was tested by having DF (for the ethogram for stress-related behaviors) and SP (for the ethogram for overall behavioral state and panting) code sets of four videos in parallel with ACVC. Cohen’s Kappa coefficient was calculated using The Observer XT. Where there was poor agreement (r<0.80), observers received further training. Values of r>0.80 were assumed to indicate strong agreement (see, for example, [8]), and once this level was attained the observers began coding videos independently. ACVC coded 77% of the videos with the ethogram for stress-related behaviors and 65% with the ethogram for overall behavioral state and panting. For each ethogram, the videos were distributed randomly between observers, with the exception that each observer coded a similar number of videos from Group Aversive and Group Reward.

##### 2.6.1.2. Cortisol analysis

Two dogs (one from School B and one from School E, both from Group Reward) did not cooperate with the saliva collection procedure and, as such, no saliva samples were extracted from them. For the remaining 90 dogs, only 23 dog owners (11 from Group Reward and 12 from Group Aversive) were able to appropriately collect six saliva samples. The samples from these subjects were selected for analysis. An additional 40 dog owners (15 from Group Reward and 25 from Group Aversive) were able to properly collect at least four saliva samples. From these 40 dogs, eight were randomly selected to have their samples analyzed (four from Group Reward and four from Group Aversive). In the end, 16 dogs from Group Aversive and 15 dogs from Group Reward had their samples selected for analysis (Schools A, C, D, E and F: n=4; School B: n=5; School G: n=6). These samples were sent to the Faculty of Sport Sciences and Physical Education of the University of Coimbra, Coimbra, Portugal, where they were assayed for cortisol concentration using standard ELISA kits (Salimetrics®).

#### 2.6.2. Phase 2 – Evaluating welfare outside the training context

For each dog, we calculated the average latency to reach the food bowl during each of the three types of test trials (NP, M, NN) as well as the average latency to reach the ‘positive’ and ‘negative’ training locations during the test phase. Afterwards, to control for individual variation in running speeds, we adjusted each dog’s test latencies by taking into account their mean latencies to get to the ‘positive’ and ‘negative’ training locations during the test phase as follows [19]:

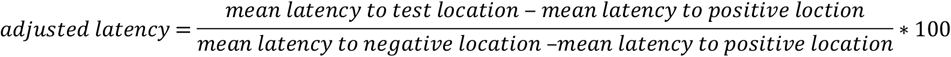

This adjusted score expresses all test latencies as a percentage of the difference between each dog’s mean latencies to the ‘positive’ and ‘negative’ locations [19].

Seventy-three dogs completed the cognitive bias task. From these 73 subjects, 14 disengaged from the task for some trials due to noise outside the test room. Thirteen disengaged for one test trial (Group Reward: one dog at location M, three dogs at location NP, and one dog at location NN; Group Aversive: five dogs at location NP, and three dogs at location NN), and one (Group Reward) for three test trials (one at each test location). For these dogs, the average latencies to the test locations were calculated from the remaining test trials. Of the remaining four dogs, one (from School G) completed the first seven test trials (at locations M,NP,NN,NP,NN,M,NN), two (one from School A and one from School E) completed the first five test trials (at locations M,NP,NN,NP,NN), and one (from School G) completed the first three test trials (at locations M,NP,NN); then they stopped cooperating with the task. Their average latencies to the test locations were calculated from these trials.

### 2.7. Statistical analyses

All statistics were conducted using the software SPSS^®^ Statistics 25.0. All data were analyzed using the Shapiro-Wilk test to check for normality. Except for the number of trials required to reach the learning criterion in the cognitive bias task, the data were not normally distributed; hence, except for the former variable, non-parametric tests were used for analysis. Namely:

- Chi-square tests were used to compare the two groups (Reward and Aversive) for dog demographics and background, and owner demographics and experience with dogs and dog training (variables collected with the questionnaire, see Appendix S3), in order to investigate whether the two groups differed in these factors.
- Kruskal-Wallis and Mann-Whitney tests were used to verify if the dog-related variables age, FCI breed group, and age of separation from the mother (which differed between Group Reward and Group Aversive, see Appendix S3) affected the dependent variables measured.
- Friedman’s ANOVA and Wilcoxon signed-rank tests were used to analyze the frequency of stress-related behaviors, the percentage of scans in the different behavioral states and the percentage of scans panting across the three training sessions, in order to examine whether there was an effect of sampling period. Where no effect of training session was found, the data for each dog were averaged across the three sessions.
- Mann-Whitney tests were used to compare the frequency of stress-related behaviors, the percentage of scans in the different behavioral states, the percentage of scans panting, the difference in cortisol concentrations and the adjusted latencies for the cognitive bias task between Group Reward and Group Aversive, in order to investigate whether the groups differed in these different indicators of welfare.
- A Wilcoxon signed-rank test was used to compare the post-training with the baseline levels of cortisol within both Group Reward and Group Aversive, with the aim of examining whether the concentration of salivary cortisol changed as a result of training for each group.
- A Mann-Whitney test was used to compare the number of training classes attended by the dogs before moving to Phase 2, to verify if these did not differ between groups.
- A t-test for independent samples was used to compare the number of trials needed to reach the learning criterion in the cognitive bias task between Group Reward and Group Aversive, in order to explore whether the groups differed in learning speed.
- A Wilcoxon signed-rank test was used to compare, in the cognitive bias task, the latency to reach the P location during test trials and during the final trial, when the bowl contained no food, to verify whether dogs were relying on olfactory or visual cues to discriminate between bowl locations.
- Spearman correlation coefficients were used to examine the correlation between the number of aversive stimuli used in the different aversive-based schools and the frequency of stress-related behaviors, the percentage of scans in the different behavioral states, the percentage of scans panting, the difference in cortisol concentrations, and the adjusted latencies in the cognitive bias task; in order to investigate whether there was a correlation between the frequency of aversive stimuli used in training and the different indicators of welfare.

With the exception of the analyses conducted to test for the effect of the sampling period on behavioral data and the analysis performed to verify if dogs were relying on olfactory or visual in the cognitive bias task, which were within-subjects, all remaining analyses were between-subjects. Two-tailed tests were used for all but correlations analyses, for which one-tailed tests were conducted. The level of significance was set at α= 0.05.

## 3. Results

### 3.1. Questionnaire

#### 3.1.1 Dog demographics and background

Concerning dog demographics, the two groups did not differ in sex and neuter status ratios, but they differed with regards to age [X^2^(3)=10.361, p=0.016] and FCI breed group [X^2^(7)=19.586, p=0.007]. As for dog background, the groups differed only in the age of separation from the mother [X^2^(7)=19.041, p=0.003, see Appendix S3 for full results]. However, statistical analysis performed for each training method group revealed that age, FCI breed group and age of separation from the mother did not affect any of the dependent variables measured in the present study (Appendix S3).

#### 3.1.2. Owner demographics, experience with dogs and dog training

Regarding owner demographics, the two groups did not differ in owner age, family household size and whether they had children, but they differed in owner sex [X^2^(1)=8.360, p=0.006]. Regarding owner experience with dogs and dog training, the groups did not differ in whether owners had attended training classes with a previous dog, but they differed in whether owners had had other dog(s) in the past [X^2^(1)=4.658, p=0.037] and in the information they relied on for choosing the dog training school [X^2^(4)=11.656, p=0.016, see Appendix S3 for full results].

### 3.2. Phase 1 – Evaluating welfare within the training context

#### 3.2.1. Behavioral data

##### 3.2.1.1. Stress-related behaviors

For each dog, we first summed all the occurrences of each stress-related behavior as defined in the ethogram for stress-related behaviors. A comparison between the two groups revealed that dogs from Group Aversive displayed significantly more stress-related behaviors than dogs from Group Reward [Group Aversive: M(±SEM)=57.06±5.98 *vs* Group Reward: M(±SEM)=10.56±1.04; U=1974.50, p<0.001; M – Mean, SEM – Standard Error of the Mean].

When analyzing each of the stress-related behaviors separately, we found that dogs from Group Aversive showed a significantly higher frequency of body turn (U=1833.50, p<0.001), move away (U=1308.50, p=0.042), crouch (U=1630.50, p<0.001), salivating (U=1211.50, p=0.019), yawn (U=1555.50, p<0.001) and lip lick (U=2004, p<0.001) than dogs from Group Reward. Additionally, there was a tendency for dogs from Group Aversive to exhibit more lying on side/back (U=1134, p=0.062), yelp (U=1174, p=0.072) and paw-lift (U=1258.50, p=0.094) behaviors. Finally, there were no differences regarding the average frequency of whine (U=1141.50, p=0.353), body shake (U=1194.50, p=0.247), and scratch (U=972, p=0.482) behaviors. Fear-related elimination was never displayed during this study (Figure 2).

**Figure 2.**
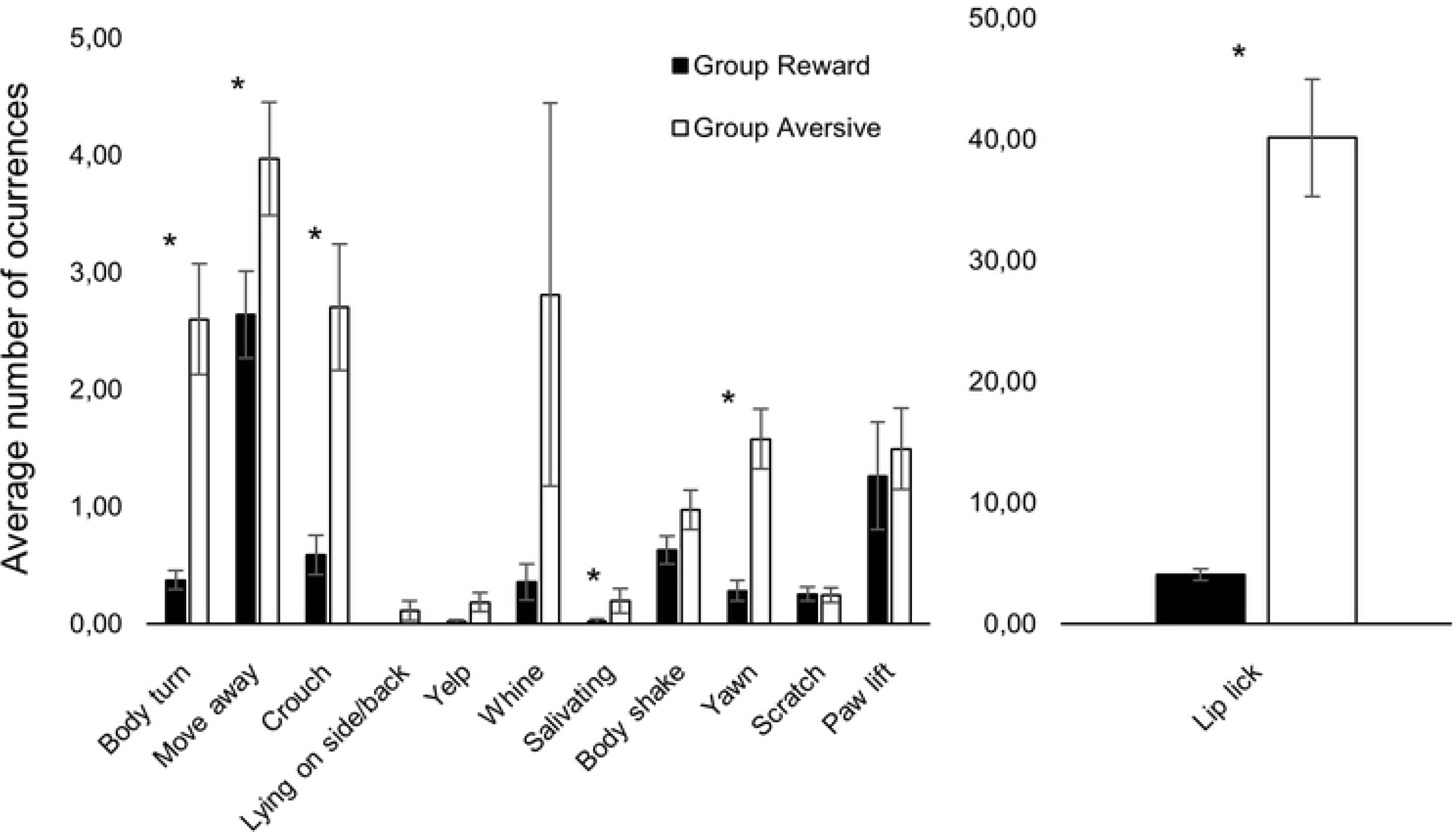
Number of occurrences of each stress-related behavior averaged across the three training sessions for Group Reward (filled bars) and Group Aversive (empty bars). Vertical bars show the SEM. * stands for statistically significant differences at α=0.05.

Because we observed a difference in the frequency of aversive stimuli used during training among the different aversive-based schools (see Appendix S3), we further analyzed the relationship between the number of stress-related behaviors displayed by the dogs and the number of aversive stimuli used in the different schools. We found a strong positive correlation between the number of aversive stimuli used and the total number of stress-related behaviors shown by the animals (Spearman correlation coefficient, r_s_=0.833, p<0.001).

Moreover, when examining the correlation between the frequency of each individual stress-related behavior and the number of aversive stimuli used by each training school, we found positive correlations for body turn (r_s_=0.653, p<0.001), move away (r_s_=0.200, p=0.028), crouch (r_s_=0.567, p<0.001), lying on side/back (r_s_=0.273, p=0.004), salivating (r_s_=0.334, p=0.001), yawn (r_s_=0.533, p<0.001), paw lift (r_s_=0.224, p=0.016) and lip lick (r_s_=0.850, p<0.001) behaviors. For the remaining behaviors there were no significant correlations (yelp: r_s_=0.107, p=0.155; whine: r_s_=0.111, p=0.147; body shake: r_s_=0.127, p=0.114; scratch: r_s_=−0.151, p=0.075).

##### 3.2.1.2. Overall behavioral state

For each training session of each dog, we calculated the percentage of scans spent in each overall behavioral state (Tense, Low, Relaxed, Excited). A Friedman’s ANOVA conducted on the percentage of scans spent in each behavioral state across the three training sessions revealed that there was a significant effect of session in the percentage of scans in the Excited state [χ^2^(2)=12.105, p=0.002], but not in the remaining states [Tense: χ^2^(2)=3.645, p=0.162; Low: χ^2^(2)=1.077, p=0.584; Relaxed: χ^2^(2)=4.244, p=0.120]. Specifically, the percentage of scans in the Excited state decreased across training sessions - Session 1: M(±SEM)=50.21±3.51, Session 2: M(±SEM)=42.94±3.59, Session 3: M(±SEM)=41.13±3.55. When analyzing the two training groups separately, we found that this pattern was statistically significant for Group Aversive [χ^2^(2)=9.490, p=0.009] but not for Group Reward [χ^2^(2)=3.674, p=0.159].

When comparing between training groups, dogs from Group Aversive showed a significantly higher percentage of scans in Tense state than dogs from Group Reward for all three training sessions [Session 1: U=1769, p<0.001; Session 2: U=1820.50, p<0.001; Session 3: U=1592, p<0.001] and a significantly higher percentage of scans in Low states for Session 1 and Session 2 [Session 1: U=1239, p=0.002; Session 2: U=1176.50, p=0.011; Session 3: U=925, p=0.211]. Additionally, dogs from Group Aversive showed a significantly lower percentage of scans in Excited [Session 1: U=296.500, p<0.001; Session 2: U=287.500, p<0.001; Session 3: U=157, p<0.001] and Relaxed [Session 1: U=628, p<0.001; Session 2: U=793, p=0.042; Session 3: U=525, p=0.001] states for all three training sessions than dogs from Group Reward (see Figure 3).

**Figure 3.**
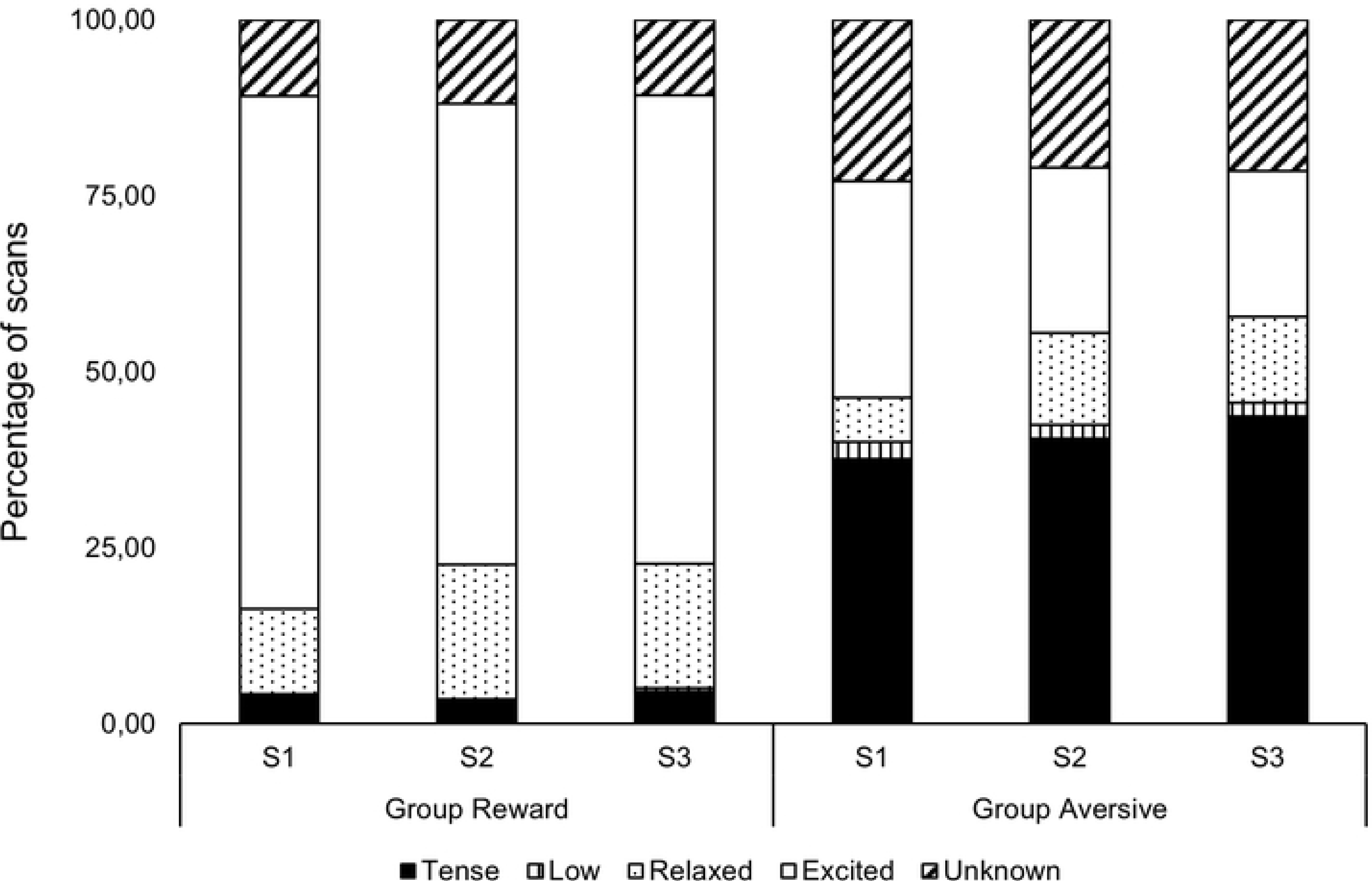
Average percentage of scans in the different behavioral states in training sessions 1 (S1), 2 (S2) and 3 (S3) for Group Reward (left) and Group Aversive (right).

When averaging the results of all three training sessions, dogs from Group Aversive showed a significantly higher percentage of scans in Tense [Group Aversive: M(±SEM)=40.50±3.07 *vs* Group Reward: M(±SEM)=4.18±0.74; U=2010.50, p<0.001] and Low [Group Aversive: M(±SEM)=2.18±0.63 *vs* Group Reward: M(±SEM)=0.16±0.12; U=1138, p<0.001] states and a lower percentage of scans in Relaxed [Group Aversive: M(±SEM)=10.03±1.95 *vs* Group Reward: M(±SEM)=16.24±2.06; U=652, p=0.002] and Excited [Group Aversive: M(±SEM)=24.92±3.27 *vs* Group Reward: M(±SEM)=68.43±2.81; U=170, p<0.001] states than dogs from Group Reward.

Finally, when examining the correlation between the percentage of scans in each behavioral state and the number of aversive stimuli used in training, we found positive correlations for Tense (r_s_=0.881, p<0.001) and Low (r_s_=0.472, p<0.001) states, and negative correlations for Relaxed (r_s_=−0.472, p<0.001) and Excited (r_s_=−0.821, p<0.001) states.

##### 3.2.1.3. Panting

For each training session of each dog, we also calculated the percentage of scans spent panting. A significant effect of training session was found for Group Reward [χ^2^(2)=7.043, p=0.030], but not for Group Aversive [χ^2^(2)=0.294, p=0.863]. However, there was no systematic increase or decrease in panting across sessions for Group Reward; instead, panting increased from Session 1 to Session 2 [Session 1: M(±SEM)= 13.93±2.91 *vs* Session 2: M(±SEM)= 20.66±3.99; T=234, p=0.054] and then decreased slightly, although not significantly, from Session 2 to Session 3 [Session 2: M(±SEM)= 20.66±3.99 *vs* Session 3: M(±SEM)=16.17±3.22; T=116.50, p=0.213]. When comparing between training groups, dogs from Group Aversive panted more than dogs from Group Reward during all training sessions (Session 1: U=1516.500, p<0.001; Session 2: U=1359, p=0.008; Session 3: U=1223.500, p=0.001, see Figure 4).

**Figure 4.**
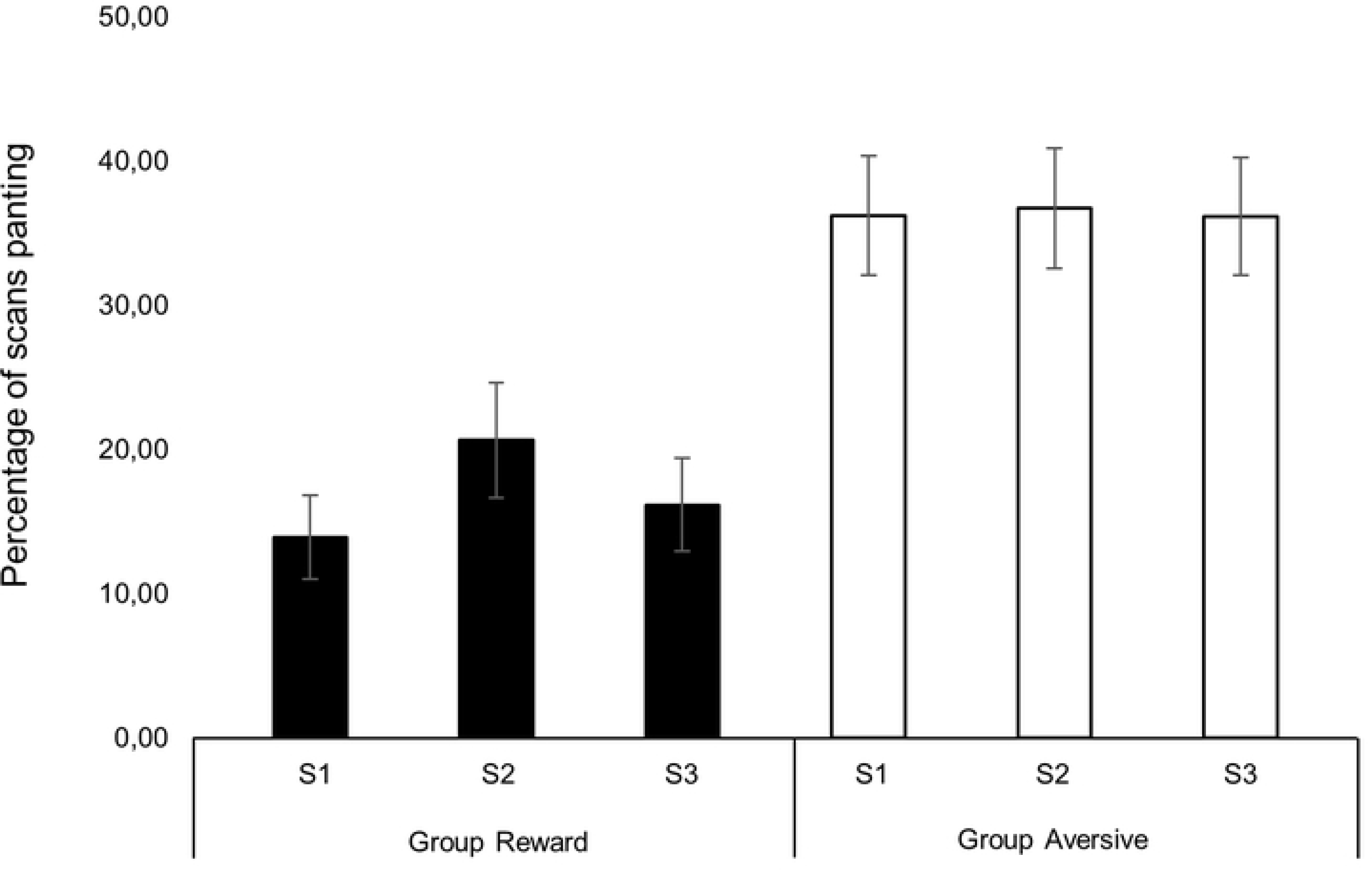
Average percentage of scans spent panting in training sessions 1 (S1), 2 (S2) and 3 (S3) for Group Reward (left) and Group Aversive (right).

When averaging the results of the three training sessions, we also found that dogs from Group Aversive panted more overall than dogs from Group Reward [Group Aversive: M(±SEM)=37.71±3.56 *vs* Group Reward: M(±SEM)=16.99±2.71; U=1540, p<0.001].

Finally, the percentage of scans spent panting was positively correlated with the number of aversive stimuli used in training (r_s_=0.407, p<0.001).

#### 3.2.2. Physiological data

In order to investigate potential changes in salivary cortisol concentration as a result of training methods, we averaged the baseline sample values (BL) and the post-training sample values (PT). Afterwards, we computed the difference between the average post-training concentration and the average baseline concentration. The average results for each training method are depicted in Figure 5. There was a significant difference between groups, with dogs from Group Aversive showing an average increase of 0.10 µg/dL in salivary cortisol concentration after training and dogs from Group Reward showing, on average, no changes in cortisol concentration (U=196, p=0.002).

**Figure 5.**
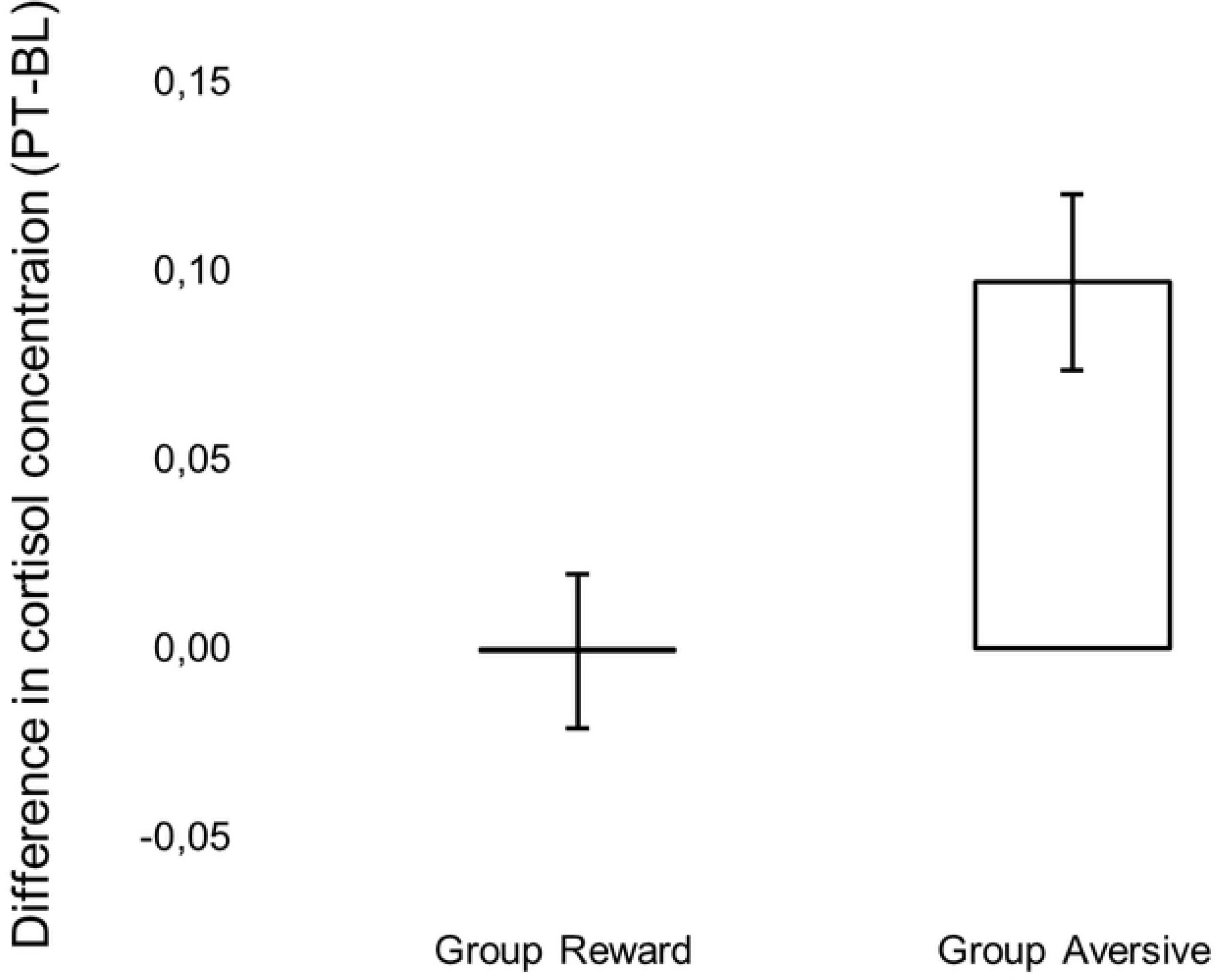
Average difference in cortisol concentration (PT: post-training average concentration, BL: baseline average concentration) for Group Aversive and Group Reward. Vertical bars show the SEM.

Additional analysis within each group revealed that the increase in cortisol concentration for Group Aversive was statistically significant [Baseline: M(±SEM)=0.14±0.01 *vs* Post-training: M(±SEM)=0.24±0.03 μ/dL; T=134, p=0.001], whereas no significant differences existed for Group Reward [Baseline: M(±SEM)=0.13±0.02 *vs* Post-training: M(±SEM)=0.13±0.02 μ/dL; T=51, p=0.609]. Moreover, whereas there were no differences between groups regarding baseline levels (U=151.50, p=0.216), the two groups differed significantly for post-training levels (U=192.50, p=0.003).

The average difference in cortisol concentrations also showed a positive correlation with the number of aversive stimuli used in training (r_s_=0.512, p<0.002).

### 3.3. Phase 2 – Evaluating welfare outside the training context

Before performing the cognitive bias task, dogs from Group Reward had attended, on average, 6.07 training classes (SEM=0.36) and dogs from Group Aversive had attended, on average, 6.66 (SEM=0.39), with no statistically significant differences observed between groups (U=1232.50, p=0.147).

#### 3.3.1. Training phase

On average, dogs took 27.14 (SEM=0.85) trials to reach the learning criterion. Dogs from the Group Reward took significantly fewer trials to reach the learning criterion than dogs from Group Aversive [Group Reward: M(±SEM)=24.80±1.26 *vs* Group Aversive: M(±SEM)=29.10±1.08, t=−2.612, p=0.011].

#### 3.3.2. Test phase

Figure 6 shows the average adjusted latencies for the two training stimuli (P, N) and the three test stimuli (NP, M, NN) for Group Reward and Group Aversive. As noted in the figure, whereas there were no significant differences between groups for either the NP (U=755.50, p=0.834) or the NN (U=874.50, p=0.154) stimuli, there was a statistically significant difference for the M stimulus, with dogs from Group Aversive taking longer to approach this bowl location than dogs from Group Reward (U=987, p=0.01).

**Figure 6.**
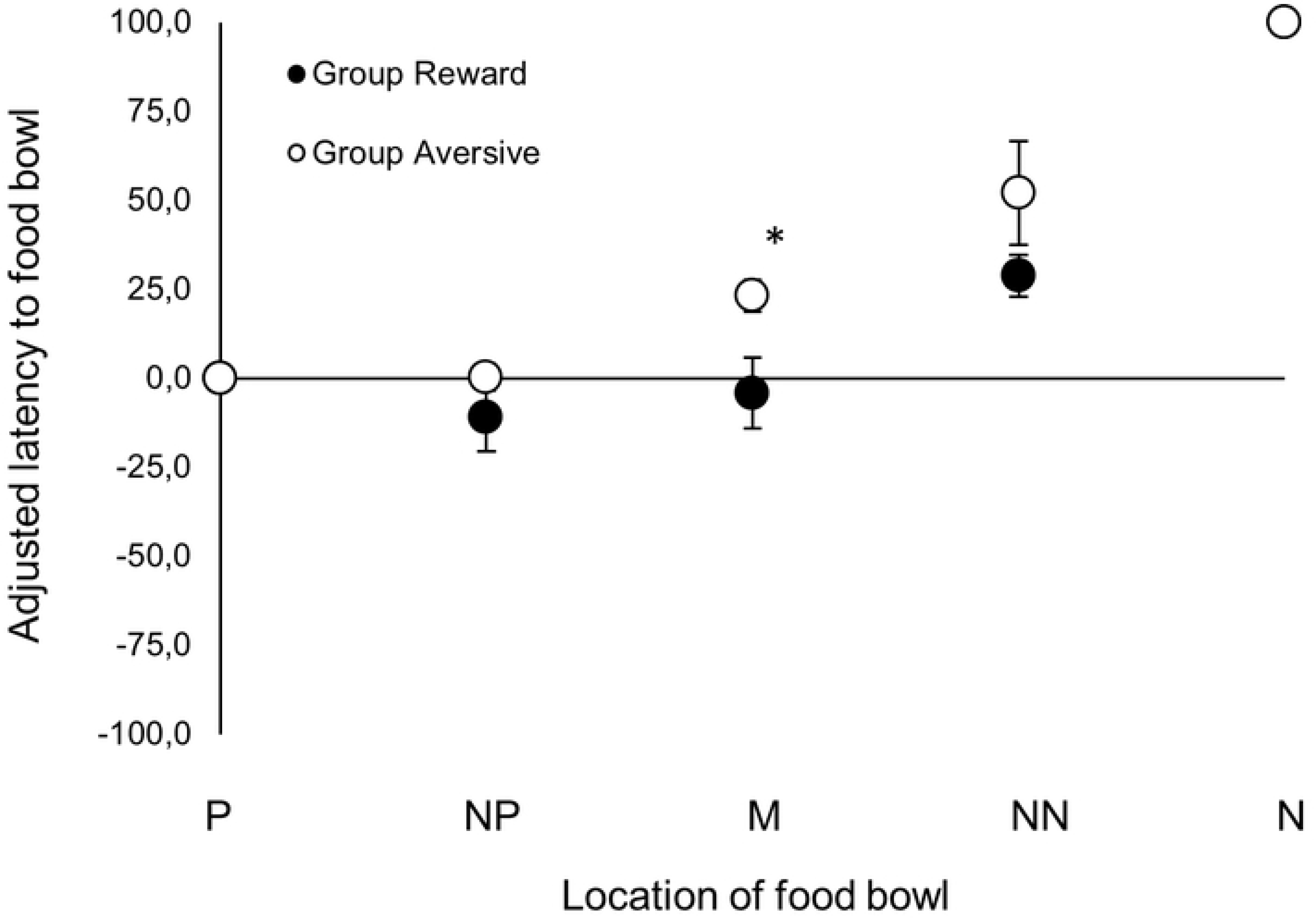
Average adjusted latency to reach the food bowl as a function of location: P - ‘positive’, NP – ‘near positive’, M – ‘middle’, NN – ‘near negative’, N – ‘negative’, for Group Reward (black circles) and Group Aversive group (white circles). Vertical bars show the SEM. * stands for statistically significant differences at α=0.017.

Additionally, we found a positive correlation between the number of aversive stimuli used in training and the adjusted latency for the M stimulus (r_s_=0.258, p=0.012). No correlations were found between the number of aversive stimuli used in training and the adjusted latencies for the NP (r_s_=−0.013, p=0.454) and the NN (r_s_=0.076, p=0.256) stimuli.

Lastly, an analysis comparing the average latency to reach the P location during test trials and the latency to reach this same location during the final trial (when the bowl contained no food) revealed no significant differences (T=1295.50, p=0.328), confirming that the dogs were not relying on olfactory or visual cues to discriminate between bowl locations.

## 4. Discussion

This was the first empirical study to systematically investigate the short- and long-term effects of aversive- and reward-based training methods on the welfare of companion dogs. We objectively classified training methods, extended the study of aversive-based methods to other techniques and tools besides shock collars, and used objective and validated measures for the assessment of both the short- (behavioral and physiological stress responses during training) and long-term welfare (cognitive bias task outside the training context) of companion dogs. Overall, our results showed that dogs trained with aversive-based methods displayed poorer indicators of both short- and long-term welfare as compared to those trained using reward-based methods.

Short-term welfare was assessed during training sessions. Here, dogs from Group Aversive spent more time in Tense and Low behavioral states as well as more time panting than dogs from Group Reward. Tense and low body postures reflect states of stress and fear in dogs (e.g., [26]), and panting has also been associated with acute stress in dogs (e.g., [8, 27]). Dogs from Group Aversive also displayed more stress-related behaviors than dogs from Group Reward. More specifically, these dogs exhibited more lip licks, yawns, body turns, moves away, crouches and instances of salivating. There was also a tendency for dogs from Group Aversive to engage in more yelping, paw-lifting and lying on side/back. In previous studies, high levels of lip licking and yawning behaviors have been consistently associated with acute stress in dogs (e.g., [9, 23]). Importantly, lip licking has been associated with stressful social situations [23]. This most likely explains the large magnitude of this behavior observed in Group Aversive, as aversive-based training methods comprise social and physical confrontation with the dog. Paw lifting, salivating and yelping have previously been interpreted as responses to pain and stress (e.g., [23, 25, 28]). The display of avoidance behaviors such as body turn, move away, crouch and lying on side/back, specifically in response to training techniques, highlights the aversive nature of the training sessions at the aversive-based schools. Notably, lying on side/back was only displayed in aversive-based schools (and mostly in School A, the school employing the highest frequency of aversive stimuli). Finally, no differences were found between groups for body shake, scratch and whine. Previous studies on dog training methods have also failed to identify significant differences regarding these behaviors [8, 9], suggesting that these behaviors may not be reliable indicators of stress, at least in the context of training. In support of this view, whining has also been associated with social solicitation, attention seeking and food begging behavior in dogs [29, 30]. It is possible that body shaking and scratching may also be associated with excitement and arousal rather than ‘negative’ stress. Hence, the present study shows a strong association between the use of aversive-based training methods and an increased frequency of stress-related behaviors during companion dog training. These results strengthen and extend the findings of previous studies on companion dogs, which suggested a positive correlation between the use of both shock collars [8] and other negative reinforcement techniques [9], and stress behaviors in the context of dog training.

With regards to physiological measures of stress, whereas Group Reward showed no differences in the concentration of salivary cortisol between baseline and post-training samples, dogs from Group Aversive exhibited a statistically significant increase in the concentration of salivary cortisol in post-training samples. Previous studies investigating cortisol levels in dogs in relation to training have yielded contradictory results. Schalke et al (2007) [31] found significant differences in the cortisol levels of three groups of laboratory dogs trained using shock collars with different degrees of shock predictability (the lower the predictability, the higher the cortisol levels). However, studies comparing aversive- and reward-based training methods have found either no significant differences or the opposite pattern. Namely, [8] and [32] found no elevation in cortisol levels after the use of shock collars and lemon-spray bark collars when compared to control groups, and [33] found that a negative punishment training method (a quitting signal) resulted in higher levels of cortisol than the use of a pinch collar (aversive-based technique). Hence, the present study is the first to report a significant increase in cortisol levels in dogs trained with aversive-based methods as compared to dogs trained with reward-based methods.

Meanwhile, the increase in cortisol levels observed in the present study (M=0.10 μg/dL) was smaller than that reported in other studies that found significant increases after dogs were exposed to aversive stimuli (0.20-0.30 μg/dL in [31] and 0.30-0.40 μg/dL in [23]). One possible explanation for this difference in magnitude may be related to the nature of the stimuli used in the different studies. Whereas the reported elevations in cortisol in [31] and [23] appeared after the presentation of non-social stimuli (shocks in [31], and shocks, sound blasts and a falling bag in [23]), the stimuli used during training in the present study where mainly of a social nature (i.e.: leash jerks, physical manipulation or yelling at the dog). Stimuli administered in a social context may be more predictable or better anticipated and, therefore, generate less acute stress responses [23]. In support of this view, [23] did not find elevations in cortisol after the presentation of social stimuli (physical restraint and opening an umbrella).

When considering long-term welfare, the present results were also in accordance with our hypothesis. Dogs from Group Aversive displayed a more ‘pessimistic’ judgment of the ‘middle’ ambiguous test location in the cognitive bias task, revealing less positive underlying affective states (and hence poorer welfare) than the dogs from Group Reward. The only other study that, to our knowledge, addressed the long-term welfare effects of training methods in dogs was performed by Christiansen et al (2001). In this study, hunting dogs were trained not to attack sheep using shock collars. No general effect of the use of shock collars on dog fear and anxiety was found one year after training took place. However, unlike the test used by [12], which was a modified version of a temperament test used by the Norwegian Kennel Club, the cognitive bias approach used in the current study is a widely established and well-validated method for evaluating animal welfare (e.g., [17, 18]). Hence, to our knowledge, this is the first study to reliably assess and report the effects of aversive- and reward-based training methods in the long-term affective states of dogs.

Additionally, dogs from Group Reward learned the cognitive bias task faster than dogs from Group Aversive. Similar findings were observed previously by [34], who found a positive correlation between the reported use of reward-based training methods and a dog’s ability to learn a novel task (touching a spoon with the nose). In another study, [35] found that dogs with high-level training experience were more successful at opening a box to obtain food than dogs that have received either none or only basic training. Although the authors reported that all subjects’ training included positive reinforcement methods, they did not specify whether positive punishment and/or negative reinforcement were used in combination. Altogether, previous research suggests that training using positive reinforcement may improve the learning ability of dogs. However, in all previous studies cited above, animals were required to perform a given behavior in order to obtain a positive reinforcer. It is unclear whether the same effect would stand if the dogs had to learn a task whose goal was, for example, to perform a behavior to escape from an unpleasant situation. It may be the case that dogs trained with positive reinforcement develop a specific ‘learning set’ [36] for tasks involving positive reinforcement, but that dogs trained with aversive-based methods perform better in tasks involving some sort of aversive stimuli. Further research is needed to clarify the relationship between training methods and learning ability in dogs.

Notably, we found that the frequency of aversive stimuli used in training (which differed among the aversive-based schools) was correlated with all the welfare indicators measured in the present study. Specifically, we found that the higher the frequency of aversive stimuli used in training, the greater the impact on the short- and on the long-term welfare of dogs. This result is in line with the findings of a previous survey study by [7], whose results showed that a higher frequency of punishment was correlated with higher anxiety and fear scores. However, even if only comparing the schools with the lowest frequency of aversive stimuli used during training (C and F) to Group Reward, differences in welfare were found (Appendix S4). Dogs trained at schools C and F showed more stress-related behaviors, spent more time in Tense and Low behavioral states and more time panting during training than dogs trained at reward-based schools, and they also took more trials to learn the cognitive bias task. However, the difference in cortisol concentration between post-training and baseline samples was only marginally significant when dogs from schools C and F were compared with dogs trained with reward-based methods, and no differences were found in the latencies to reach the ambiguous stimuli in the cognitive bias task. This seems to suggest that, although dogs trained in ‘least aversive’ schools show poorer indicators of welfare than dogs trained in reward-based schools, these are not as poor as those of dogs trained in ‘most aversive’ schools. Notably, schools C and F also used positive reinforcement frequently during their training sessions alongside aversive-based methods, which was rarely the case in schools A and D. Nonetheless, caution is required when interpreting these results due to the reduced sample size for analysis of only two training schools. Future studies should further compare the effects on dog welfare of reward-based training and what can be called ‘balanced’ training, or the use of reward-based methods alongside aversive-based methods. Despite the fact that the results of the present study show that reward-based methods are better for dog welfare, the efficacy of training should also be examined. Presently, there is a lack of scientific evidence regarding the efficacy of different training methods (see also [3]), and thus further research in the topic is required. If reward-based methods are, as the current results show, better for dog welfare than aversive-based methods, and also prove to be more effective or equally effective to aversive-based methods, there is no doubt that owners and dog professionals should use reward-based training practices. If, on the other hand, aversive-based methods prove to be more effective, we would advise using aversive stimuli as infrequently as possible during training, and use them in combination with reward-based techniques, due to the implications for dog welfare.

Ultimately, some limitations of the present study must be considered. Firstly, because this was an empirical rather than an experimental study, we cannot infer a true causal relationship between training methods and dog welfare. To do so would require a randomized control trial. Because we did not randomly allocate dogs to the two treatments (training methods), we cannot discard the possibility that dogs enrolling training at aversive-based schools had higher stress levels *a priori*, or that there are other significant differences between dog-owner pairs that lead some owners to choose an aversive-based school and others to choose a reward-based school. The fact that the baseline levels of cortisol did not differ between groups weakens the former possibility. Regarding the latter, however, we did indeed find differences between groups in owner sex, previous experience with dogs and in the information they relied on for choosing the dog training school. However, conducting an experimental study would raise ethical concerns (but see [8]), as previous studies have already suggested an association between the use of aversive-based methods and indicators of stress in dogs (see [3] for a review), as well as with the quality of dog-owner attachment [37]. Secondly, we need to consider the possibility for a volunteer bias and hence any generalization of the present results must take this in account.

## 5. Conclusions

Our results show that companion dogs trained using aversive-based methods experienced poorer welfare as compared to companion dogs trained using reward-based methods, at both the short- and the long-term level. Specifically, dogs attending schools using aversive-based methods displayed more stress-related behaviors and body postures during training, higher elevations in cortisol levels after training, and were more ‘pessimistic’ in a cognitive bias task than dogs attending schools using reward-based methods. Moreover, we found that the higher the frequency of aversive stimuli used in training, the greater the impact on the short- and the long-term welfare of dogs. To our knowledge, this is the first comprehensive and systematic study to evaluate and report the effects of dog training methods on companion dog welfare. Critically, our study points to the fact that the welfare of companion dogs trained with aversive-based methods appears to be at risk.

## Acknowledgements

We are grateful, first and foremost, to all dogs and their owners who participated in this study; without them this research would never have been possible. A very special acknowledgment to the dog training schools and their trainers that opened their doors for our participant recruitment and data collection.

We would also like to thank Joana Guilherme-Fernandes for the support provided in the development of the setup for the cognitive bias task, for helping with data collection and especially for all the invaluable discussions during study planning and data interpretation. A special acknowledgment also for Margarida Lencastre and Flávia Canastra, who also helped in data collection. Finally, we want to thank Jennifer Barrett for input given during data collection and analysis.

The current research study was supported by FCT - Fundação Portuguesa para a Ciência e Tecnologia (Fellowship SFRH/BPD/111509/2015) and UFAW – Universities Federation for Animal Welfare (Grant 14-16/17). Stefania Pastur was supported by PIPOL - Regione Friuli Venezia Giulia.

## Supporting information captions

Appendix S1. Information on training schools and training methods.

Table S1a. Characterization of the training schools by types of behaviors trained, training sites and class structure.

Table S1b. Definition of the aversive-based operant conditioning procedures used to classify the dog training schools as aversive-based or reward-based. The schools were classified as aversive-based if they used some sort of positive punishment and/or negative reinforcement and as reward-based if they did not use any of these techniques.

Table S1c. Frequency (mean ± standard deviation) of positive punishment and negative reinforcement used during the four training sessions videotaped at the dog training schools. Schools A, C, D and F were classified as aversive-based and Schools B, E and G were classified as reward-based.

Appendix S2. Ethograms used for the analysis of the video recordings of dog training sessions.

Table S2a. Ethogram for stress-related behaviors.

Table S2b. Ethogram for overall behavioral state and panting.

Appendix S3. Questionnaire data and relationship with the dependent variables measured in the present study.

Table S3a. Variables obtained from the questionnaire (dog demographics and background, and owner demographics and experience with dogs and dog training). Chi-square tests were used to compare the two groups (Reward and Aversive).

Table S3b. Statistical results for the effects of the dog-related variables that differed between the Reward and the Aversive groups (age, FCI breed group and age of separation from the mother) on the different dependent variables measured in the current study. The variable age comprised five categories (< 6 months; 6-11 months; 1-3 years; 4-7 years; >7 years), the FCI breed group variable comprised eleven (Mixed breed; Sheepdogs and Cattledogs, except Swiss Cattledogs; Pinscher and Schnauzer – Molossoid and Swiss Mountain and Cattledogs; Terriers; Daschunds; Spitz and primitive types; Scent hounds and related breeds; Pointing dogs; Retrievers, Flushing Dogs and Water Dogs; Companion and Toy Dogs; Sight Hounds), and the variable age of separation from the mother comprised nine [less than 1 month; 1 – 1.5 months (inclusive); 1.5 – 2 months (inclusive); 2 – 2.5 months (inclusive); 2.5 – 3 months (inclusive); 3 – 4 months (inclusive); 4 – 5 months (inclusive); more than 5 months, don’t know]. Kruskal-Wallis and Mann-Whitney tests were used to compare more or less than two categories, respectively. Significant differences were found for the frequency of paw-lift and percentage of scans in Tense state in Group Aversive and for the percentage of scans panting in Group Reward; however, post-hoc pairwise comparisons with adjusted Bonferroni’s correction revealed no differences between the different categories.

Appendix S4. Statistical results for the comparison of the different welfare indicators between Group Reward and the two schools from Group Aversive that used the lowest frequency of aversive stimuli during training (C and F).

Appendix S5. Raw data underlying all the analyzes performed in the current research paper.

## References

1. Reid P. Learning in dogs. In: Jensen P, editor. The Behavioural Biology of Dogs. CABI; 2007. pp. 120–144.

2. Reisner I. The learning dog: A discussion of training methods. In: Serpell J, editor. The Domestic Dog: Its Evolution, Behavior and Interactions with People, 2nd Edition. Cambridge University Press; 2017. pp. 211–226.

3. Guilherme-Fernandes J, Olsson IAS, Vieira de Castro AC. Do aversive-based training methods actually compromise dog welfare?: A literature review. Appl Anim Behav Sci. 2017; 196, 1–12.

4. Hiby EF, Rooney NJ, Bradshaw JWS. Dog training methods: their use, effectiveness and interaction with behaviour and welfare. Anim Welf. 2004; 13, 63–69.

5. Blackwell EJ, Twells C, Seawright A, Casey RA. The relationship between training methods and the occurrence of behavior problems, as reported by owners, in a population of domestic dogs. J Vet Behav. 2008; 3: 207–217.

6. Herron ME, Shofer FS, Reisner IR. Survey of the use and outcome of confrontational and non-confrontational training methods in client-owned dogs showing undesired behaviors. Appl Anim Behav Sci. 2009; 117, 47–54.

7. Arhant C, Bubna-Littitz H, Bartles A, Futschik A, Troxler J. Behaviour of smaller and larger dogs: effects of training methods, inconsistency of owner behavior and level of engagement in activities with the dog. Appl Anim Behav Sci. 2010; 123: 131–142.

8. Cooper JJ, Cracknell N, Hardiman J, Wright H, Mills D. The welfare consequences and efficacy of training pet dogs with remote electronic training collars in comparison to reward based training. PLoS One. 2014; 9, e102722.

9. Deldalle S, Gaunet F. Effects of 2 training methods on stress-related behaviors of the dog (Canis familiaris) and on the dog-owner relationship. J Vet Behav. 2014; 9, 58–65.

10. Companion Animal Welfare Council, 2012. The Use of Electric Pulse Training Aids (EPTAs) in Companion Animals. Available from: http://eprints.lincoln.ac.uk/14640/1/CAWC%20ecollar%20report.pdf (Accessed 12 August 2019).

11. Haverbeke A, Laporte B, Depiereux E, Giffroy JM, Diederich C. Training methods of military dog handlers and their effects on the team’s performances. Appl Anim Behav Sci. 2008; 113, 110–122.

12. Christiansen FO, Bakken M, Braastad BO. Behavioural changes and aversive conditioning in hunting dogs by the second-year confrontation with domestic sheep. Appl Anim Behav Sci. 2001; 72, 131–143.

13. Wiepkema PR, Koolhaas JM. Stress and animal welfare. Anim Welfare. 1993; 2: 195–218.

14. Conrad CD. A critical review of chronic stress effects on spatial learning and memory. Prog Neuro-Psychoph. 2010; 34: 742–755.

15. Destrez A, Deiss V, Lévy F, Calandreau L, Lee C, Chaillou-Sagon E. Chronic stress induces pessimistic-like judgment and learning deficits in sheep. Appl Anim Behav Sci. 2013; 148: 28–36.

16. Harding EJ, Paul ES, Mendl M. Animal behaviour: cognitive bias and affective state. Nature. 2004; 427: 312–312. PMID: 14737158.

17. Boissy A, Erhard HW. How studying interaction between animal emotion, cognition, and personality can contribute to improve farm animal welfare. In: Grandin T, Deesing MJ, editors. Genetics and the Behavior of Domestic Animals, 2nd Edition. Academic Press. 2014. pp. 81–113.

18. Mendl M, Burman OHP, Parker RMA, Paul ES. Cognitive bias as an indicator of animal emotion and welfare: emerging evidence and underlying mechanisms. Appl Anim Behav Sci. 2009; 118, 161–181.

19. Mendl M., Brooks J, Basse C, Burman O, Paul E, Blackwell E, Casey R. Dogs showing separation-related behaviour exhibit a ‘pessimistic’ cognitive bias’. Curr Biol. 2010; 20, R839–R840.

20. Barker SB, Wolen AR. The benefits of human-companion animal interaction: a review. J Vet Med Educ. 2008; 35(4): 487–495.

21. Crawford EK, Worsham NL, Swinehart ER. Benefits derived from companion animals, and the use of the term “attachment”. Anthrozoos. 2006; 19 (2), 98–112

22. Dreschel NA, Granger DA. Methods of collection for salivary cortisol measurement in dogs. Horm Behav. 2009; 55(1): 163–188.

23. Beerda B, Schilder MBH, van Hooff JARAM, de Vries HW, Mol JA. Behavioural, saliva cortisol and heart rate responses to different types of stimuli in dogs. Appl Anim Behav Sci. 1998; 58, 365–381.

24. Kobelt A, Hemsworth PH, Barnett JL, Butler KL. Sources of sampling variation in saliva cortisol in dogs. Res Vet Sci. 2003; 75: 157–161.

25. Schilder MBH, van der Borg JAM. Training dogs with help of the shock collar: short and long term behavioural effects. Appl Anim Behav Sci. 2004; 85: 319–334.

26. Beerda B, Schilder MBH, van Hooff JARAM, de Vries HW. Manifestations of chronic and acute stress in dogs. Appl Anim Behav Sci. 1997; 52:307–319.

27. Voith VL, Borchelt PL. Fears and phobias in companion animals. In: Voith VL, Borchelt PL, editors. Readings in companion animal behaviour. Trenton, NJ: Veterinary Learning Systems; 1996. pp. 140–152.

28. De Palma C, Viggiano E, Barillari E, Palme R, Dufour AB, Fantini C, Natoli E. Evaluating the temperament in shelter dogs. Behaviour. 2005; 142: 1307–1328.

29. Mills DS, Beral A, Lawson S. Attention seeking behavior in dogs–what owners love and loathe! J Vet Behav. 2010; 5: 60.

30. Pongrácz P, Molnár C, Miklósi A, Csanyi V. Human listeners are able to classify dog (Canis familiaris) barks recorded in different situations. J Comp Psy. 2005; 119: 136–144.

31. Schalke E, Stichnoth J, Ott S, Jones-Baade R. Clinical signs caused by the use of electric training collars on dogs in everyday life situations. Appl Anim Behav Sci. 2007; 105: 369–380.

32. Steiss JE, Schaffer C, Ahmad HA, Voith VL. Evaluation of plasma cortisol levels and behavior in dogs wearing bark control collars. Appl Anim Behav Sci. 2007; 106, 96–106.

33. Salgirli Y, Schalke E, Hackbarth H. Comparison of learning effects and stress between 3 different training methods (electronic training collar, pinch collar and quitting signal) in Belgian Malinois Police Dogs. Rev Méd Vét. 2012; 163, 530–535.

34. Rooney NJ, Cowan S. Training methods and owner-dog interactions: links with dog behaviour and learning ability. Appl Anim Behav Sci. 2011; 132, 169–177.

35. Marshall-Pescini S, Valsecchi P, Petak I, Accorsi PA, Prato-Previde E. Does training make you smarter? The effects of training on dogs’ performance (Canis familiaris) in a problem solving task. Behav Process. 2008; 78: 449–454.

36. Harlow HF. The formation of learning sets. Psychol Rev. 1949; 56: 51–65.

37. Vieira de Castro AC, Barrett J, de Sousa L, Olsson IAS. Carrots versus stick: The relationship between training methods and dog-owner bond. Appl Anim Behav Sci. 2019; 219: 104831.

